# Are all waterholes equal from a lion’s view? Exploring the role of prey abundance and catchability in waterhole visitation patterns in a savannah ecosystem

**DOI:** 10.1101/2025.04.18.649579

**Authors:** Romain Dejeante, Andrew J. Loveridge, David W. Macdonald, Daphine Madhlamoto, Simon Chamaillé-Jammes, Marion Valeix

## Abstract

Prey abundance and catchability shape the spatial ecology of predators. Predators can select habitats where prey are more abundant to maximize encounter rate with prey or habitats where prey are more catchable to maximize prey capture. These hypotheses are commonly referred to as prey-abundance and prey-catchability hypotheses. Although these hypotheses are often tested at the landscape scale, little is known about how between-patch variations in prey abundance and catchability determine the space use of predators. In many savannah ecosystems, large herbivores aggregate around waterholes, which become hotspots of prey and their selection by predators is classically interpreted as supporting the prey-abundance hypothesis. Here, we investigated whether between-waterhole variations in prey abundance and catchability influence the frequency and duration of lion visits to waterholes, testing the prey-abundance and prey-catchability hypotheses at the resource-patch scale. We combined datasets on (1) lion movements recorded from GPS collars deployed on 20 adult males and 16 adult females between 2002 and 2015, (2) prey abundance evaluated from long-term, regular monitoring of waterholes and (3) prey catchability evaluated from remote-sensing satellite imagery of vegetation cover around waterholes in Hwange National Park (Zimbabwe). Lions did not use all waterholes in their territory equally: there was a high variability in the frequency and duration of visits. Surprisingly, between-waterhole variations in prey abundance and catchability only slightly explained these variations in frequency – and even less in duration – of lion visits to waterholes. Yet, the frequency of lion visits to waterholes decreased with the number of waterholes within their territory, and male lions more frequently visited the waterholes surrounded by more open habitats. We discuss the limits of our work, but also the ecological mechanisms that may explain these findings. First, lions and their prey are involved in a ‘shell-game’ that leads them to adopt unpredictable movement strategies. Second, lions have only access to a limited number of waterholes amongst which to distribute their hunting effort. Lastly, lions visit waterholes not only to hunt but also to interact with social mates and competitors. This work challenges the implicit assumption that all waterholes are the same from a lion’s view and calls for further studies investigating the drivers of the variability in lion visits at the resource-patch scale.

## Introduction

The study of how predator-prey interactions shape the spatial behaviour of both predators and prey has a long history in the ecological literature (Creel et al., 2005; Fortin et al., 2005; Gaynor et al., 2019; Gilliam & Fraser, 1987; Palmer et al., 2022). Predators and prey are involved in a ‘shell game’ in which they endlessly adjust their behaviour in response to one another (Mitchell & Lima, 2002; Sih, 2005; Simon et al., 2019) with prey attempting to avoid predators and predators attempting to be unpredictable for prey (Patin et al., 2020). From a predation perspective, predators are classically expected to use habitats more intensively where prey are more abundant, seeking to maximize the encounter rate with prey (i.e., “prey-abundance hypothesis”) (Davidson et al., 2012; Kittle et al., 2017; Zabihi-Seissan et al., 2022). They are also expected to use habitats more intensively where prey are more catchable, seeking to maximize prey capture (i.e., “prey-catchability hypothesis”) (Balme et al., 2007; Hopcraft et al., 2005; Zabihi-Seissan et al., 2022). However, these hypotheses are not mutually exclusive and predators should adjust their spatial behaviour to variations in both prey abundance and catchability, i.e., prey availability (Smith et al., 2020). Despite this recognition, the roles of prey abundance and prey catchability on predator spatial behaviour have largely been examined separately, limiting our understanding of their relative importance or their possible interactive influence on spatial ecology of predators. For example, do predators consistently favour habitats where prey are more catchable, or do they seek to maximize prey-catchability solely where prey are locally scarce?

Unfortunately, ecologists often lack knowledge on prey abundance and catchability at the resource-patch scale to investigate the intricacies between these two measures. Instead, prey abundance and catchability are commonly evaluated at the landscape scale by considering some habitat types as a proxy for prey abundance, and others as a proxy for prey catchability (see for example Balme et al. 2007; Kauffman et al. 2007; Smith et al. 2020). In savannah ecosystems, waterholes and open grasslands are intensively used by large herbivores (Courbin et al., 2019; Smit et al., 2007; Valeix, Loveridge, et al., 2009) and their selection by predators is interpreted as supporting the prey-abundance hypothesis (Davidson et al., 2012; Valeix et al. 2010). Conversely, tall grasslands, woody vegetation, thickets or rocky outcrops offer numerous ambush sites for stalk-and-ambush predators and their selection by such predators is interpreted as supporting the prey-catchability hypothesis (Davidson et al., 2012; Davies et al., 2016; Hopcraft et al., 2005). However, comparing a predator’s selection for several habitat types does not allow ecologists to address how predators respond to between-patch variation in prey abundance and catchability. Indeed, prey abundance and catchability likely vary spatially from one site to another – see for example Chamaillé-Jammes et al. (2016) for between-waterhole variations in herbivore distribution – and predators may seek to maximize differently prey capture or prey encounter rate depending on these local variations.

An important gap in our understanding of the spatial ecology of predators is therefore predator responses to between-patch variations in prey abundance and catchability. Here, to explore the prey-abundance and prey-catchability hypotheses at the between-patch scale, we investigated the spatial responses of African lions (*Panthera leo*) to between-waterhole variations in prey abundance and catchability in a savannah ecosystem, Hwange National Park, Zimbabwe, where the role of waterholes on the spatial ecology of lions has been extensively studied.

In this ecosystem, waterholes are known to attract large herbivores, particularly in the dry season (Valeix, Loveridge, et al., 2009; Valeix, 2011), and thus can be considered as prey hotspots, and ultimately key hunting sites for lions (Davidson et al., 2012; Valeix et al., 2010; Valeix, Fritz, et al., 2009). They even shape lion territory size (Loveridge et al., 2009; Valeix et al., 2012) and the way lions use their territory (Dejeante, Loveridge, Macdonald, Madhlamoto, Chamaillé-Jammes, et al., 2025). However, most of these studies did not consider heterogeneity in the characteristics of waterholes. The abundance of herbivores varies, spatially, from one waterhole to another, and between-waterhole variations in prey abundance are consistent over years and substantially explain the spatial structure of herbivores communities (Chamaillé-Jammes et al., 2016). Further, whereas some waterholes are located in large open grasslands, others are located in woodlands or bushlands, influencing the catchability of herbivores near these waterholes, as lions are stalk-and-ambush hunters expected to have a higher hunting success in environments where vegetation can provide concealment (Davies et al., 2016; Funston et al., 2001; Schaller, 1972). How lions adjust their space use to between-waterhole variations in both prey abundance and catchability remains unknown.

The intensity of use of a location – although commonly evaluated in movement ecology, for example to estimate an animal’s home range or an animal’s habitat selection – has the limitation of aggregating how frequently and for how long an individual visits a location, which restricts our understanding of the animal’s spatial ecology (Bastille-Rousseau et al., 2024; Benhamou & Riotte-Lambert, 2012). Here, we therefore investigated how between-waterhole differences in prey abundance and catchability influence the frequencies and durations of lion visits to the waterholes available in their territory. In particular, following the *prey-abundance hypothesis*, we predicted that lions would more frequently use waterholes with a higher prey abundance and for a longer duration. However, such a response may depend on the average prey abundance within lion territories: for example, lions may spend more time close to a waterhole with a higher prey abundance when prey are generally scarce within their territory, but may not do so if prey are abundant throughout their territory *(functional response in prey-abundance hypothesis)*. Following the *prey-catchability hypothesis*, because lions are predators that commonly hunt by stalking and ambushing their prey (Funston et al., 2001; Schaller, 1972), we predicted that lions would more frequently use those waterholes with a higher prey catchability (i.e., less open vegetation or short distance between vegetation cover and waterhole bank) and for a longer duration. Similarly, we tested whether lions respond more strongly to between-waterhole variations in prey catchability when prey were scarce in their territories *(functional response in prey-catchability hypothesis)*. Finally, because lions may adjust how often they visit a waterhole and the time they spend close to this waterhole to both the local variations in prey abundance and catchability simultaneously, we also predicted that they should maximize the use of waterholes with both high prey abundance and high prey catchability *(functional response in both prey-abundance and prey-catchability hypotheses)*.

## Methods

### Study area

The study was conducted in the north-eastern area of Hwange National Park, Zimbabwe. The park covers 14,600 km² of semi-arid savannah. In this study, we used data collected during the late dry season, from July to mid-October, when natural rain-fed pans are dry and surface water remains available only in artificial permanent waterholes in which underground water is pumped (Chamaillé-Jammes et al., 2007). A GIS layer of all artificial permanent waterholes that retain water throughout the dry season was available.

### Lion visits to waterholes

We investigated lion visits to waterholes from movement data collected from 20 adult male lions and 16 adult female lions between 2002 and 2015. Lions were monitored with GPS collars, collecting hourly or two-hourly locations. As lions were commonly equipped over more than one year, we used animal-year as our sampling unit, resulting in 38 animal-year units for males, and 35 animal-year units for females. Because prey-abundance data (see the following section) was only available during the dry season, only movement data from the late dry season, between July and mid-October, were analysed. Lion handling and care protocols were consistent with guidelines provided in the ‘Code of Practice for Biologists using Animals’, Department of Biology, University of Oxford and approved by University of Oxford, Biomedical Sciences, Animal Welfare and Ethical Review Body. Lion handling procedures were carried out by project staff trained and certified by the Zimbabwe Veterinary Association, Wildlife Group (Certificate numbers: Davidson: 09/03; Elliot: 9/10; Hunt: A11/04, 005/09, 5/14; Loveridge: 6/2000, 20/36 (2007), 6/14; Stapelkamp: 34/2008) in accordance with Statutory Instrument 409 of 1999 (Clause 21A to 21J) amending the Regulations of 1975 to the Dangerous Drugs Act, Zimbabwe.

For each animal-year, we evaluated the frequency and the duration of visits to each waterhole of a lion territory using the R package *recurse* (Bracis et al., 2018) to count, respectively, the number of times a collared lion entered a circular buffer around a waterhole – over 100 days – and to measure the time spent within this buffer per visit. Following Bracis et al., (2018), the entrance and exit times for each visit were calculated by linear interpolation between the trajectory locations inside and outside the buffer. We performed our analyses using both a 1km and a 2km radius around each waterhole. These distances are commonly employed to investigate the behaviour of lions in the vicinity of waterholes in the study ecosystem (Courbin et al., 2016; Davidson et al., 2013; Dejeante et al., 2024; Valeix, Fritz, et al., 2009; Valeix et al., 2010). We used the two distance thresholds to test the sensitivity of our results to the chosen value. Results did not differ qualitatively, and we present results obtained with the 1km radius threshold in the main text. Results from the analyses with the 2km radius threshold are available in the Supplementary Information (Tables S1 and S2). Because the number of days of tracking during the dry season varied from one animal-year to another – from 60 to 105 days – we re-calculated the number of visits to a waterhole over a period of 100 days using a proportional relationship. This allowed us to compare the number of visits from one animal-year to another.

We estimated the frequency and the duration of lion visits to each waterhole within their territory, as delineated by the minimum convex polygon created from 100% of lion’s movement data collected during the late dry season. Although using kernel-based methods is commonly a more revealing approach to delineate an animal’s home range (Börger et al., 2006; Nilsen et al., 2008), here we used a minimum convex polygon to consider as available all waterholes potentially reachable by an individual, even those rarely or never visited. Contrary to kernel-based methods, drawing a minimum convex polygon does not allow us to measure differences in the intensity of use. However, these relative differences in the intensity of use of each available waterhole within a lion territory were explored from the differences in the frequency and duration of visits. In the following analyses, we seek to explain differences in the frequency and duration of lion visits to each waterhole of their territory by between-waterhole variations in prey abundance and catchability.

### Prey abundance data

To evaluate prey abundance at each waterhole, we used a long-term, regular monitoring of waterholes conducted by the Wildlife Environment Zimbabwe (WEZ) since 1972. Every year, at the end of the dry season, the WEZ records the animals coming to drink to each waterhole of the park over a 24h period (all waterholes with water are monitored). This monitoring provides information on the total number of herds and the average herd size that visited a waterhole for all large herbivore species. Importantly, Chamaillé-Jammes et al. (2016) showed that the spatial structure of the herbivore community is mostly explained by between-waterhole variations and that differences in prey abundance between waterholes are consistent over years. Although the park has a wide diversity of large herbivores, here, we focused only on the main prey species of lions (Davidson et al., 2013): African buffalo (*Syncerus caffer*), plains zebra (*Equus quagga*), blue wildebeest (*Connochaetes taurinus*), roan antelope (*Hippotragus equines*), sable antelope (*Hippotragus niger*), greater kudu (*Tragelaphus strepsiceros*) and eland (*Taurotragus oryx*). For species other than buffalo, because encounter rates between predators and prey are better predicted by prey herd density than by prey individual density (Fryxell et al., 2007), and because their herd size does not show a high variability in Hwange National Park (Supplementary Information Table S3), we used the total number of herds drinking at a waterhole (pooling all species) as a proxy for medium-sized prey abundance at this waterhole. Hunting medium-sized prey is generally different from hunting larger prey, such as buffalo. Large prey are indeed harder and more dangerous to catch but provide more food (Funston et al., 1998, 2001). Besides, buffaloes in the study ecosystem are mostly found in a few very large groups (hundreds of individuals; with groups as large as ∼1000 individuals) that have fixed and large territories (Wielgus, 2020). Hence although they regularly visit the waterholes in their territory, the likelihood of not observing them during a 24h period is high. As buffaloes are a particularly important prey species for lions and given this low probability of detection, we chose not to aggregate buffaloes and other prey, but to average buffalo data over a 5-year window, to maximize the chance of the large buffalo herd detection at this waterhole. As the buffalo herds range over stable spatial areas over time, we believe that using a 5-year window allowed us to minimize the detection bias.

Variables related to medium-sized prey or buffalo abundance were centred and scaled per animal-year territory (hereafter ‘Herd’ and ‘Buffalo’ variables) to test whether, within a territory, lions visit more waterholes with higher abundance in herds of medium-sized prey or in buffalos (‘prey-abundance hypothesis’). In addition, we also calculated the mean herd and buffalo abundance per animal-year within each lion territory (‘Herd Territory’ and ‘Buffalo Territory’) and then centred and scaled this variable across all animal-year territories. Adding an interaction term between the ‘Herd’ and ‘Herd Territory’ variables allowed us to test whether lions visit more waterholes with higher herd abundance specifically when their territory is poor or rich in herds (‘functional response in prey-abundance hypothesis’).

### Prey catchability data

In Hwange National Park, the vegetation is dominated by bushlands and woodlands interspersed with patches of grasslands surrounding waterholes (Arraut et al., 2018). In this analysis, we considered two scales. First, we used the distance between the waterhole bank and the vegetation cover (measured using Google Earth Pro © for each cardinal direction (North, South, East and West) and then averaging these 4 distances for each waterhole) as a proxy for the fine-scale prey catchability at a waterhole (Figure 1; ‘Distance to cover’ variable). Second, we used the mean vegetation openness within the 1km surrounding each waterhole as a proxy for the large-scale prey catchability around a waterhole (‘vegetation openness’ variable). To do so, we used the 30-m resolution vegetation map produced by Arraut et al. (2018) to calculate, for each pixel, the proportion of open vegetation (category ‘grassland’ and ‘open bushlands’ in the original map) in a radius of 250 m. We then averaged the proportion of open vegetation within a radius of 1km around each waterhole. For each metric, we assumed that prey in more open areas (i.e., high distance between the waterhole and the vegetation cover or higher proportion of open vegetation) are less catchable because of the better visibility conditions to detected an approaching lion – and that lions should therefore more frequently visit (or over longer periods) the waterholes surrounded by less open areas if prey catchability drives their movement decisions at the patch-scale.

**Figure 1.**
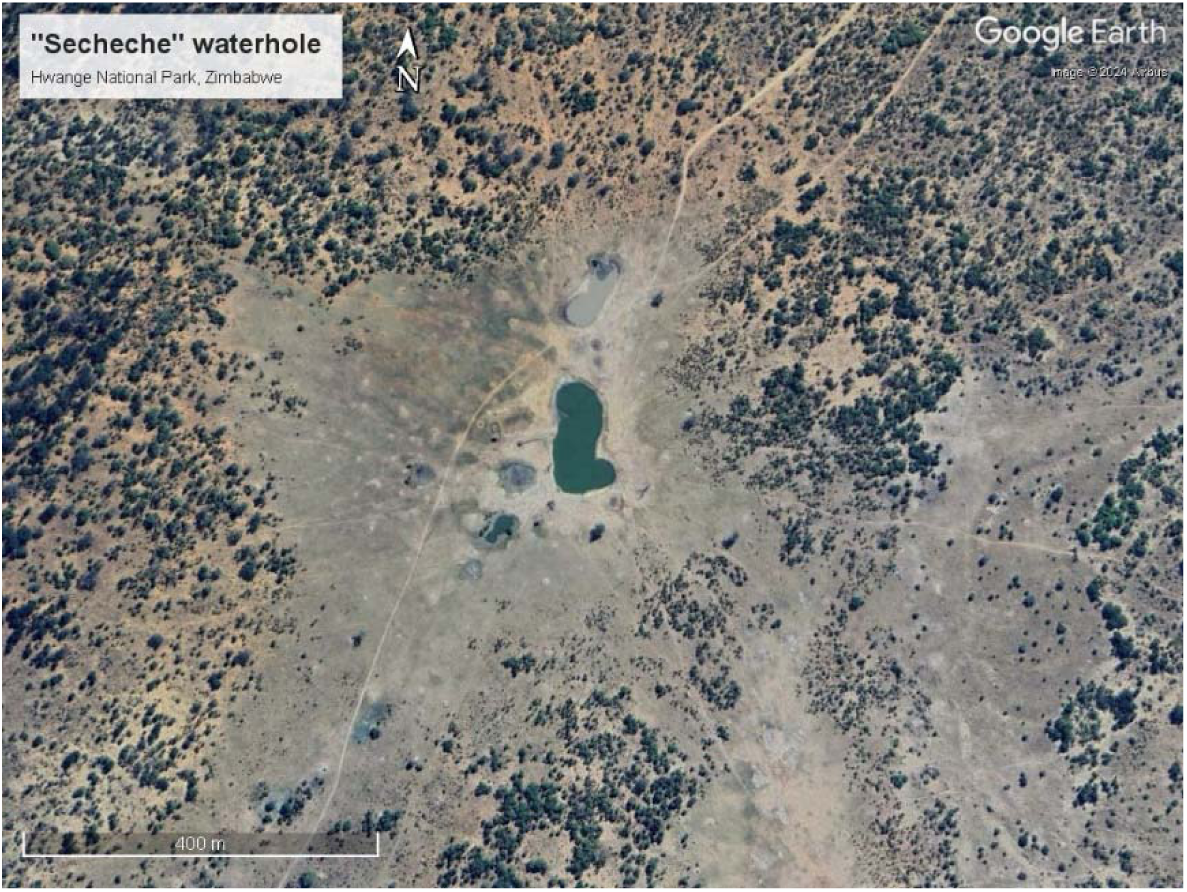
A patch of grassland surrounding a waterhole in Hwange National Park, Zimbabwe. Photo Google Earth Pro ©

### Data analysis

We investigated how between-waterhole variations in prey abundance and catchability influenced the differences in the frequency and duration of lion visits to the different waterholes of their territory. Since the frequency and duration of lion visits to waterholes exhibited overdispersion in Poisson models, we used negative binomial regressions to model these response variables. For each waterhole within a lion territory, we tested the influence of the explanatory variables related to prey abundance and prey catchability on the frequency or on the duration of visits to the corresponding waterhole by the corresponding lion. Because of the structure of the data, all models included a random intercept with animal-year identity. Model sets are presented in Tables 1 and 2 and are grouped according to the tested hypotheses (see Introduction). Because the frequency of visits to a waterhole and the duration of these visits may not be related solely to prey availability but also to the number of waterholes within the lion territory, we controlled for this effect by including the number of waterholes within the territory of an animal-year as a fixed effect in all tested models.

**Table 1.**
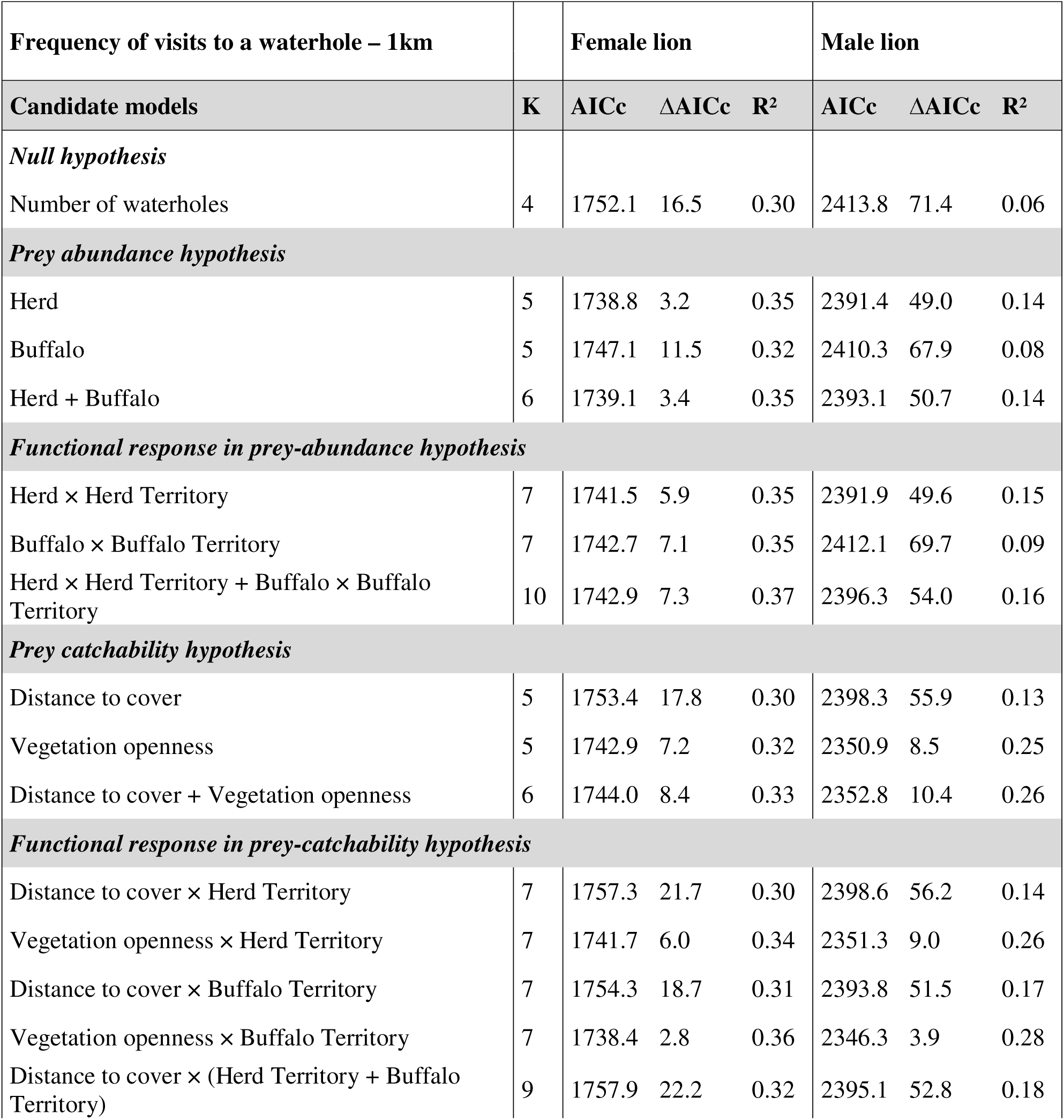
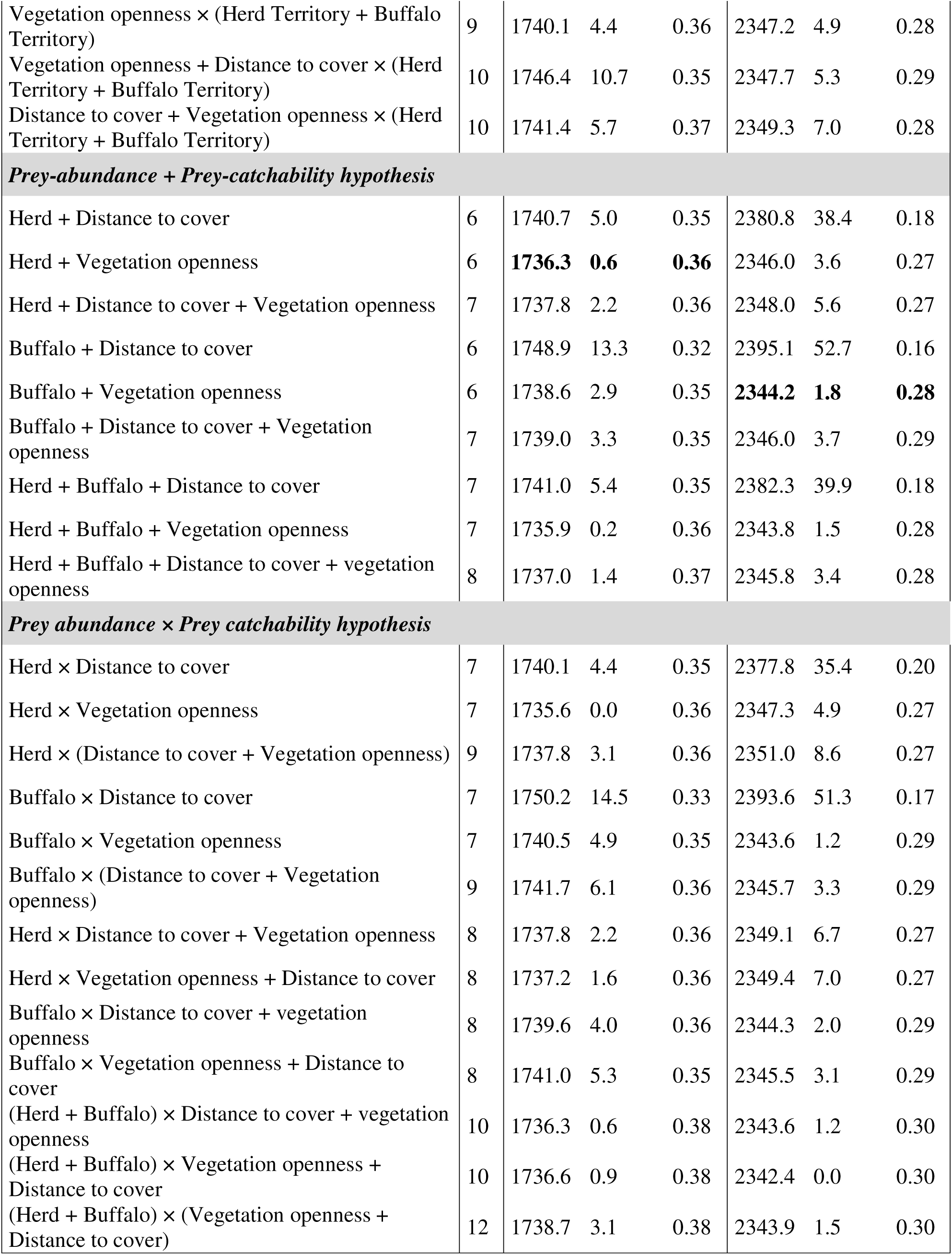
Summary of the candidate models explaining the variability of the frequency (i.e., the number of visits over 100 days) of lion visits to a waterhole. All models included the number of waterholes within the territory of each animal-year and a random effect with the identity of each animal-year. The most parsimonious models (ΔAICc < 2 and lowest number of parameters *K*) are shown in bold. For these models, we used a distance threshold of 1km to consider that lions visited a waterhole.

**Table 2.** Summary of the candidate models explaining the variability of the duration of lion visits to a waterhole. All models included the number of waterholes within the territory of each animal-year and a random effect with the identity of each animal-year. The most parsimonious model (ΔAICc < 2 and lowest number of parameters *K*) is shown in bold. For these models, we used a distance threshold of 1km to consider that lions visited a waterhole.

| **Duration of visit to a waterhole – 1km** |  | **Female lion** |  |  | **Male lion** |  |  |
| --- | --- | --- | --- | --- | --- | --- | --- |
| **Candidate models** | **K** | **AICc** | **$\Delta\text{AICc}$** | **R<sup>2</sup>** | **AICc** | **$\Delta\text{AICc}$** | **R<sup>2</sup>** |
| ***Null hypothesis*** |  |  |  |  |  |  |  |
| *Number of waterholes* | 4 | **4026.6** | **0.0** | **0.03** | 5020.0 | 6.1 | 0.06 |
| ***Prey abundance hypothesis*** |  |  |  |  |  |  |  |
| Herd | 5 | 4028.5 | 1.9 | 0.03 | 5019.1 | 5.1 | 0.07 |
| Buffalo | 5 | 4028.4 | 1.8 | 0.03 | 5018.0 | 4.1 | 0.07 |
| Herd + Buffalo | 6 | 4030.0 | 3.4 | 0.03 | 5018.8 | 4.9 | 0.08 |
| ***Functional response in prey-abundance hypothesis*** |  |  |  |  |  |  |  |
| Herd × Herd Territory | 7 | 4030.1 | 3.5 | 0.04 | 5020.4 | 6.5 | 0.08 |
| Buffalo × Buffalo Territory | 7 | 4031.8 | 5.2 | 0.03 | 5021.5 | 7.6 | 0.07 |
| Herd × Herd Territory + Buffalo × Buffalo Territory | 10 | 4035.6 | 9.0 | 0.05 | 5024.6 | 10.7 | 0.09 |
| ***Prey catchability hypothesis*** |  |  |  |  |  |  |  |
| Distance to cover | 5 | 4028.2 | 1.6 | 0.03 | 5020.3 | 6.3 | 0.06 |
| Vegetation openness | 5 | 4028.0 | 1.4 | 0.03 | **5015.8** | **1.9** | **0.08** |
| Distance to cover + Vegetation openness | 6 | 4030.0 | 3.4 | 0.03 | 5017.9 | 4.0 | 0.08 |
| ***Functional response in prey-catchability hypothesis*** |  |  |  |  |  |  |  |
| Distance to cover × Herd Territory | 7 | 4030.3 | 3.8 | 0.04 | 5022.1 | 8.2 | 0.07 |
| Vegetation openness × Herd Territory | 7 | 4028.4 | 1.8 | 0.05 | 5017.5 | 3.5 | 0.08 |
| Distance to cover × Buffalo Territory | 7 | 4031.7 | 5.1 | 0.03 | 5021.6 | 7.7 | 0.07 |
| Vegetation openness × Buffalo Territory | 7 | 4027.5 | 0.9 | 0.05 | 5016.0 | 2.1 | 0.09 |
| Distance to cover × (Herd Territory + Buffalo Territory) | 9 | 4034.2 | 7.6 | 0.04 | 5023.3 | 9.4 | 0.08 |
| Vegetation openness × (Herd Territory + Buffalo Territory) | 9 | 4030.0 | 3.4 | 0.06 | 5016.2 | 2.3 | 0.10 |
| Vegetation openness + Distance to cover × (Herd Territory + Buffalo Territory) | 10 | 4036.2 | 9.6 | 0.04 | 5021.1 | 7.2 | 0.10 |
| Distance to cover + Vegetation openness × (Herd Territory + Buffalo Territory) | 10 | 4032.1 | 5.5 | 0.06 | 5018.3 | 4.4 | 0.10 |
| ***Prey abundance + Prey catchability hypothesis*** |  |  |  |  |  |  |  |
| Herd + Distance to cover | 6 | 4030.2 | 3.6 | 0.03 | 5019.6 | 5.6 | 0.07 |
| Herd + Vegetation openness | 6 | 4030.0 | 3.4 | 0.03 | 5016.8 | 2.8 | 0.08 |
| Herd + Distance to cover + Vegetation openness | 7 | 4032.0 | 5.4 | 0.03 | 5018.8 | 4.9 | 0.08 |
| Buffalo + Distance to cover | 6 | 4029.9 | 3.3 | 0.03 | 5018.6 | 4.6 | 0.08 |
| Buffalo + Vegetation openness | 6 | 4029.7 | 3.1 | 0.03 | 5013.9 | 0.0 | 0.09 |
| Buffalo + Distance to cover + Vegetation openness | 7 | 4031.7 | 5.1 | 0.03 | 5016.0 | 2.1 | 0.09 |
| Herd + Buffalo + Distance to cover | 7 | 4031.5 | 4.9 | 0.03 | 5019.5 | 5.5 | 0.08 |
| Herd + Buffalo + Vegetation openness | 7 | 4031.6 | 5.0 | 0.03 | 5015.8 | 1.9 | 0.09 |
| Herd + Buffalo + Distance to cover + vegetation openness | 8 | 4033.6 | 7.0 | 0.03 | 5017.9 | 4.0 | 0.09 |
| ***Prey abundance × Prey catchability hypothesis*** |  |  |  |  |  |  |  |
| Herd × Distance to cover | 7 | 4032.1 | 5.5 | 0.03 | 5021.2 | 7.3 | 0.07 |
| Herd × vegetation openness | 7 | 4030.1 | 3.5 | 0.04 | 5018.8 | 4.9 | 0.08 |
| Herd × (Distance to cover + Vegetation openness) | 9 | 4034.2 | 7.6 | 0.04 | 5021.6 | 7.6 | 0.08 |
| Buffalo × Distance to cover | 7 | 4031.8 | 5.2 | 0.03 | 5019.5 | 5.6 | 0.08 |
| Buffalo × Vegetation openness | 7 | 4030.2 | 3.6 | 0.04 | 5016.0 | 2.1 | 0.09 |
| Buffalo × (Distance to cover + Vegetation openness) | 9 | 4033.2 | 6.6 | 0.04 | 5018.7 | 4.8 | 0.09 |
| Herd × Distance to cover + Vegetation openness | 8 | 4034.0 | 7.4 | 0.03 | 5019.9 | 5.9 | 0.08 |
| Herd × Vegetation openness + Distance to cover | 8 | 4032.1 | 5.5 | 0.04 | 5020.9 | 7.0 | 0.08 |
| Buffalo × Distance to cover + Vegetation openness | 8 | 4033.6 | 7.0 | 0.03 | 5016.8 | 2.9 | 0.09 |
| Buffalo × Vegetation openness + Distance to cover | 8 | 4032.2 | 5.6 | 0.04 | 5018.1 | 4.1 | 0.09 |
| (Herd + Buffalo) × Distance to cover + Vegetation openness | 10 | 4036.9 | 10.3 | 0.04 | 5020.3 | 6.4 | 0.10 |
| (Herd + Buffalo) × Vegetation openness + Distance to cover | 10 | 4035.5 | 8.9 | 0.04 | 5022.1 | 8.1 | 0.09 |
| (Herd + Buffalo) × (Vegetation openness + Distance to cover) | 12 | 4038.7 | 12.1 | 0.05 | 5023.5 | 9.6 | 0.10 |

To determine the most parsimonious model explaining the differences of frequency and duration of lion visits to a waterhole, we used the Akaike information criterion corrected for small sample size (AICc) (Burnham & Anderson, 2004). In the following, we present results obtained from the most parsimonious models, i.e., with both a ΔAICc < 2 and the lowest number of explanatory variables (Arnold, 2010). Selected models show no multicollinearity between predictors, as indicated by variance inflation factors (VIF) lower than 2. Analyses were conducted using the R package *glmmTMB* (Brooks et al., 2017). We used the R package *performance* (Lüdecke et al., 2021) to measure the VIF scores and the goodness-of-fit measure of the models as estimated by the marginal pseudo-R² for generalized linear mixed models (Nakagawa et al., 2017).

## Results

### Influence of prey abundance and catchability on the frequency of lion visits to a waterhole

On average, lions visited a given waterhole in their territory 8 times (for female) or 9 times (for males) over a 100-day period of the late dry season, but a very large variability was observed, and some waterholes could be visited by the same lion more than 50 times (Figure 2.a-b). For male lions, the most parsimonious model explaining the variability of the frequency of visits to a waterhole included the number of waterholes in the territory, the number of buffalo herds visiting the waterhole and the vegetation openness in the surroundings of the waterhole (Table 1; *Buffalo + Vegetation openness*). Details of model coefficients and errors are available in the Supplementary Information (Table S4). Male lions more frequently visited waterholes surrounded by open vegetation and those more frequently used by buffalo herds, relative to other waterholes available in their territories (Figure 3). Unsurprisingly, they also less frequently visited each waterhole when their territory encompassed a larger number of waterholes. Overall, this model explained 28% of the variability in the number of visits to a waterhole. The vegetation openness was the main variable explaining this variability. Indeed, the null model – that solely included the number of waterholes – explained only 6% of this variability and only 8% when also accounting for buffalo abundance against 25% when accounting for the vegetation openness (and not accounting for buffalo abundance) (Table 1).

**Figure 2.**
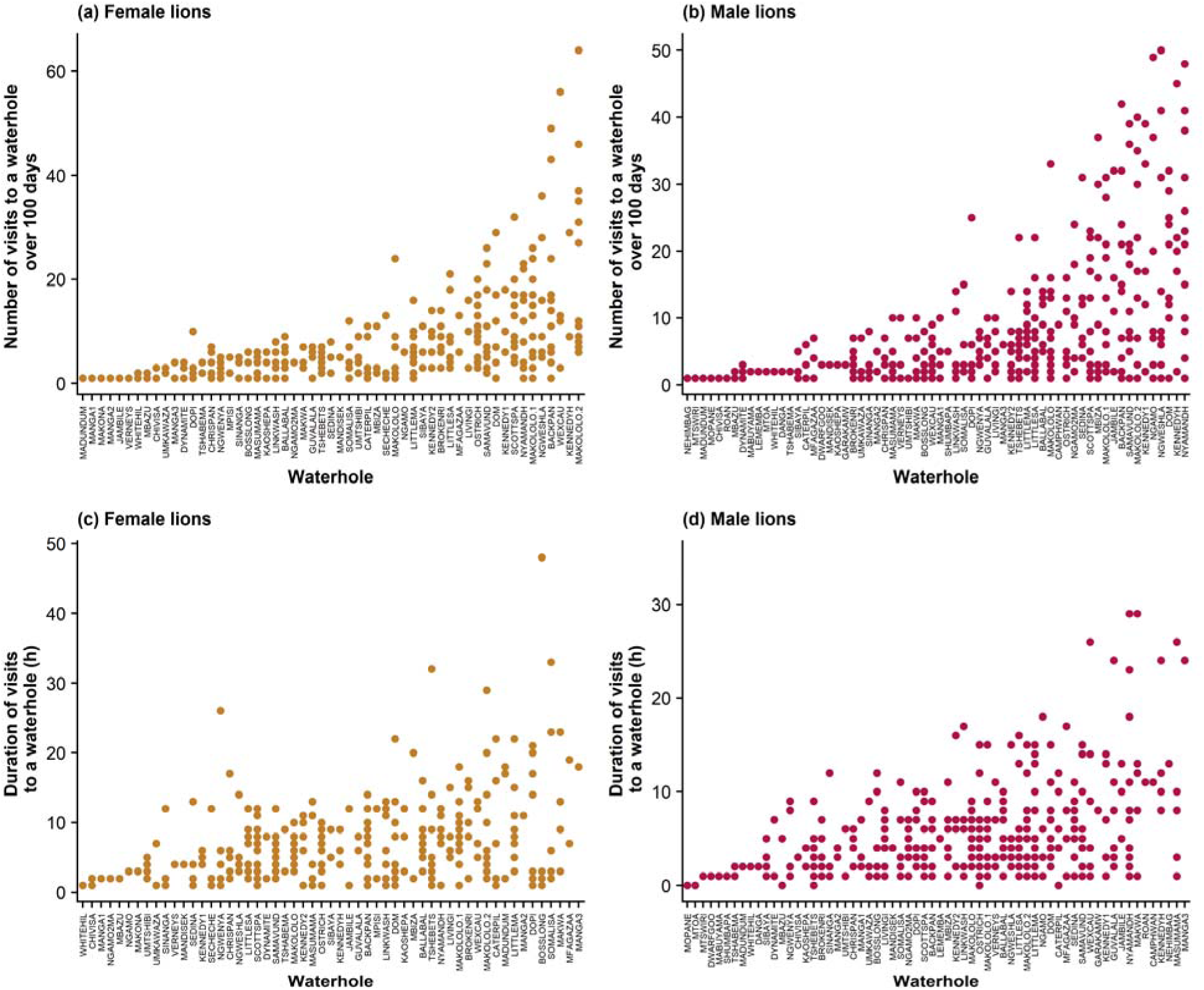
Distribution of the frequencies and durations of lion visits to a specific waterhole during the dry season. We used a 1km distance threshold to define lion visits to a waterhole. For male lions, one visit to a waterhole lasting 76h is not shown in the plot. Waterholes along the x-axis are ordered by the average frequency of visits by female lions (a) and male lions (b) or by the average duration of visits by female lions (c) or male lions (d). The x-axis shows the name of each waterhole. Each dot shows the frequency or duration of visits to the corresponding waterhole for one animal-year.

**Figure 3.**
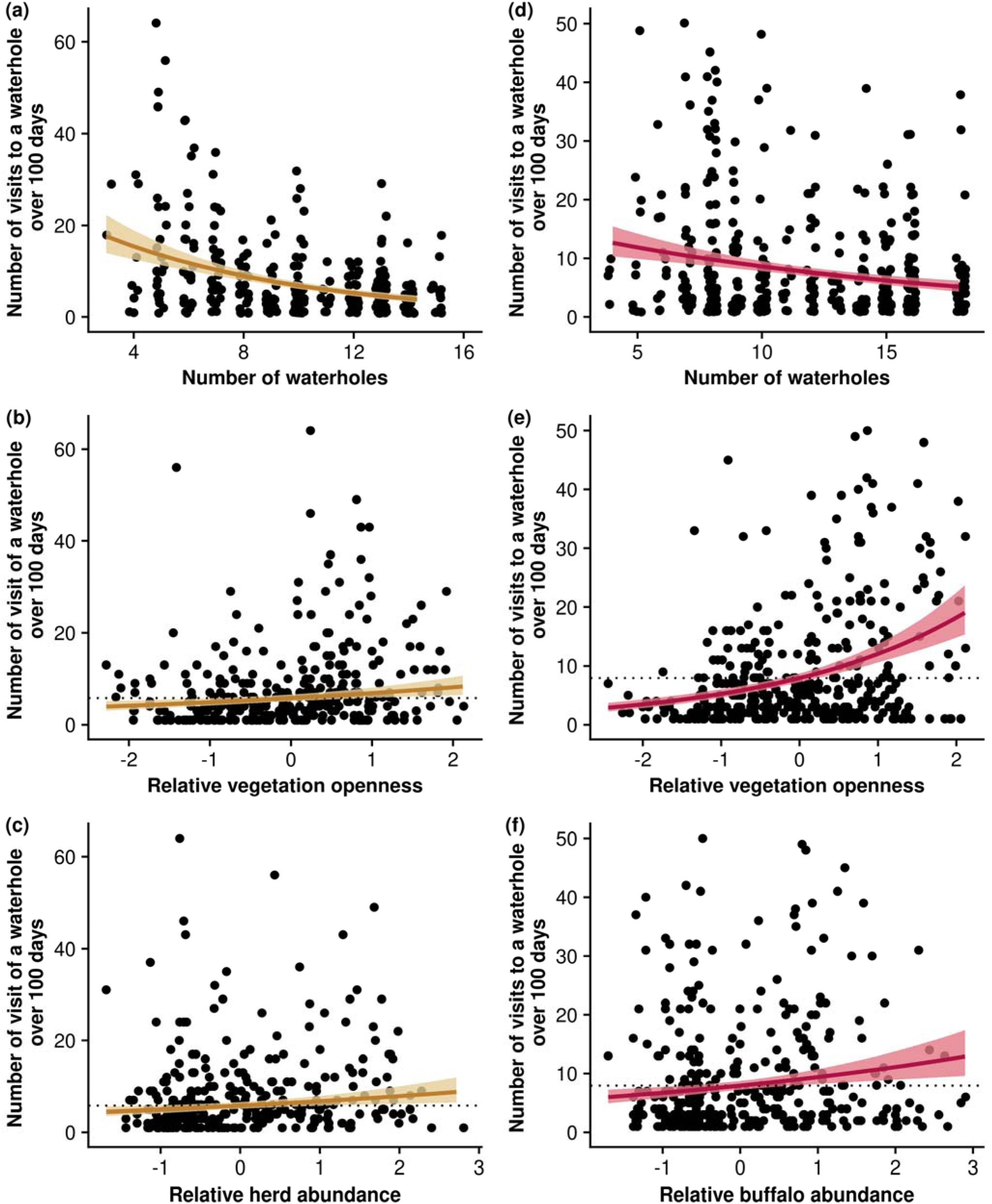
Influence of prey abundance and catchability on the frequency of visits to a specific waterhole by female (yellow) and male (red) lions. Ribbon extremities show 95% confidence interval, whereas lines show the mean value of the response variable. Predictions are made from models using a 1km distance threshold to consider that lions visit a waterhole. The dotted horizontal line shows the frequency of visits to a waterhole expected from the null hypothesis (i.e., the frequency of lion visits to a waterhole solely depends on the number of waterholes within their territory). Prey-related metrics are centred and scaled per animal-year territory to compare the relative abundance and the relative catchability of prey from one waterhole to another waterhole available in the animal-year territory.

For female lions, the most parsimonious model explaining the variability of the frequency of visits to a waterhole included the number of waterholes in the territory, the number of herds of medium-sized prey and the vegetation openness in the surroundings of the waterhole (Table 1; *Herd + Vegetation openness*). Details of model coefficients and errors are available in the Supplementary Information (Table S5). Female lions more frequently visited waterholes surrounded by open vegetation and those more frequently used by herds of medium-sized prey (Figure 3). As for male lions, females also less frequently visited each waterhole when the number of waterholes in their territory increased. Although this model explained 36% of the variability in the frequency of visits to a waterhole by female lions, the metrics related to prey abundance and catchability explained only a small amount of this variability. Indeed, the null model – that included the number of waterholes only – explained 30% of total variability. This result suggests that the vegetation openness and the number of medium-sized prey herds only slightly affect the frequency of visits to a waterhole by female lions.

### Influence of prey abundance and catchability on the duration of lion visits to a waterhole

On average, lions visited a given waterhole in their territory for 7 hours (for females) or 6 hours (for males) per visit, but with high variability between waterholes, as some visits could last more than 24 hours (Figure 2.c-d). The relationship between durations and frequencies of lion visits to a given waterhole of their territories is available in the Supplementary Information (Figure S1). For both male and female lions, all models tested only slightly explained the variability in the duration of visits to a waterhole. For female lions, the most parsimonious model included solely the number of waterholes in the territory (Table 2; *null hypothesis*) and only explained 3% of the variability in the duration of visits (Figure 4). Details of model coefficients and errors are available in the Supplementary Information (Table S6). For male lions, the most parsimonious model, which included the number of waterholes in the territory and the vegetation openness in the surroundings of the waterhole (Table 2; *Vegetation openness*) – only explained 9% of this variability. Details of model coefficients and errors are available in the Supplementary Information (Table S7). Although male lions stayed longer at waterholes where vegetation openness was greater than in surrounding areas (Figure 4), the low pseudo-R² of this model suggests that this variable only slightly affect the duration of visits.

**Figure 4.**
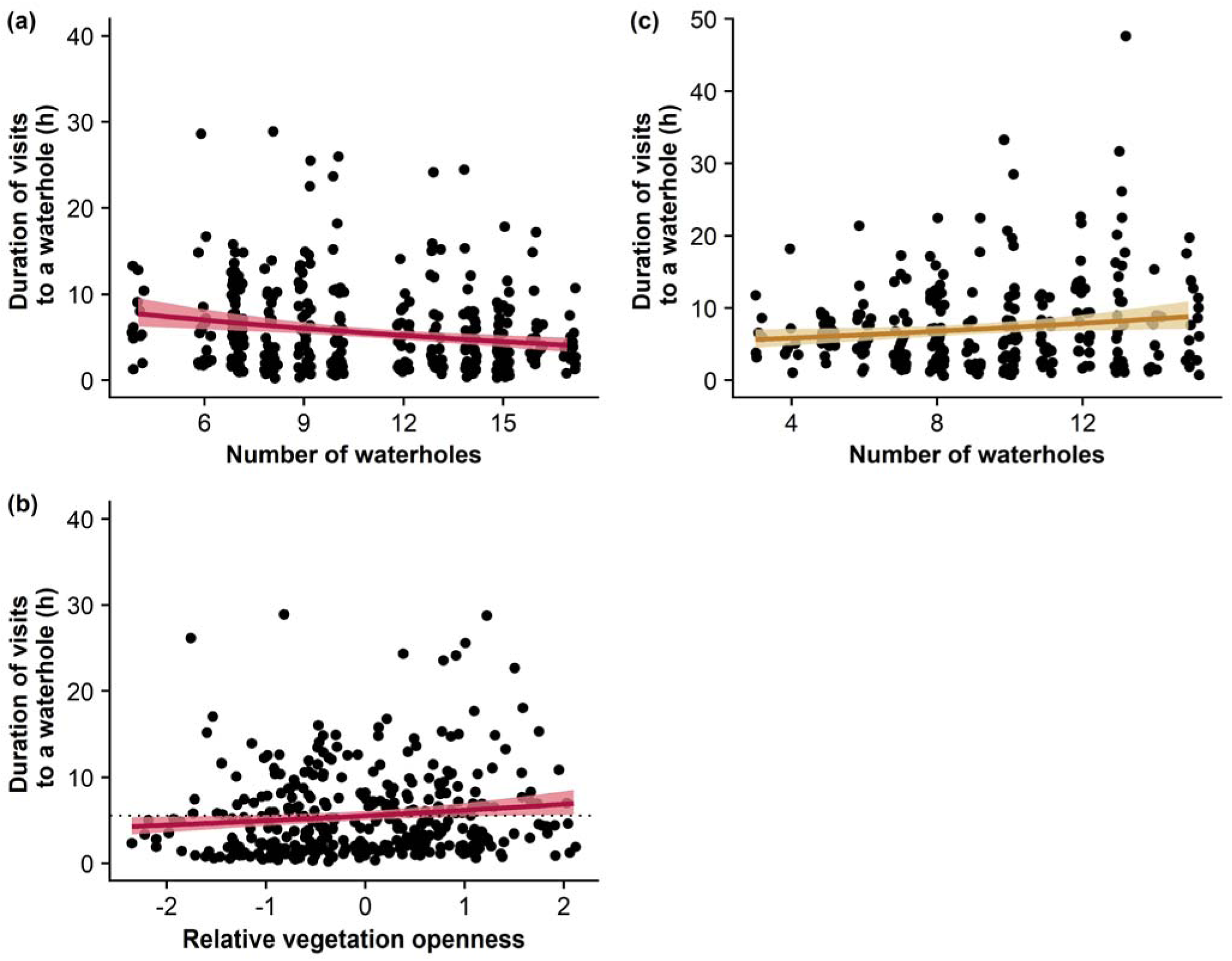
Influence of the number of waterholes within lions territories and of the relative vegetation openness in the surroundings of a waterhole on the durations of visits to a specific waterhole by female (yellow) and male (red) lions. Ribbon extremities show 95% confidence interval, whereas lines show the mean value of the response variable. Predictions are made from models using a 1km distance threshold to consider that lions visit a waterhole. The dotted horizontal line shows the duration of visits to a waterhole expected from the null hypothesis (i.e., the duration of lion visits to a waterhole solely depends on the number of waterholes within their territory). For male lions, one visit to a waterhole lasting 76h is not shown in the plot.

## Discussion

In savannah ecosystems, waterhole areas are key foraging patches for predators and their prey (Davidson et al., 2012; Mosser et al., 2015; Valeix et al., 2010; Valeix, Fritz, et al., 2009; Valeix, Loveridge, et al., 2009). However, whether all waterholes are similarly attractive from a predator’s viewpoint or whether predators favour some waterhole areas rather than others remains unknown. Here, we investigated how between-waterhole variations in prey abundance and catchability influenced the frequency and duration of lion visits to a waterhole. Surprisingly, we found that the prey-related variables considered in this work only slightly affected the frequency – and even less the duration – of visits to a waterhole by male and female lions. Our main findings are that (1) lions do not equally use (different frequencies and durations of visits) the waterholes in their territory, (2) the duration of visits to waterholes by lions is not influenced by the associated prey abundance and vegetation openness, (3) lions of both sexes tend to more frequently visit waterholes in their territory when there are fewer waterholes overall, (4) prey abundance (as measured in this work) does not affect the frequency of visits to a waterhole by lions, and (5) male lions tend to more often visit waterholes surrounded by open vegetation. The ecological mechanisms and the limits of our work that may explain these findings are discussed below.

### A dynamic predator-prey space race

In ecosystems with defined prey hotspots, predators are expected to employ a ‘leap-frogging’ behaviour (Sih, 2005) whereby their spatial patterns are shaped by these prey hotspots. This has been documented for wolves (Kittle et al., 2017) and lions (Kittle et al., 2016; for Hwange: Dejeante, Loveridge, Macdonald, Madhlamoto, Chamaillé-Jammes, et al., 2025). Although predators can rely on cues of higher prey abundance or higher prey catchability to move between these prey hotspots (Balme et al., 2007; Davidson et al., 2012; Hopcraft et al., 2005), predator and prey are involved in a ‘shell game’ in which they endlessly adjust their behaviour in relation to one another to minimize the predictability of their presence (Gaynor et al., 2019; Mitchell, 2009; Mitchell & Lima, 2002). Indeed, if predators are too predictable, prey can proactively increase their vigilance when foraging close to well-known ambush site or can avoid foraging nearby. Because of this predator-prey space race, prey can seek to remain unpredictable to predators and can be reluctant to use some waterholes despite their importance in the landscape or use them during the day when the risk of predation is lower (Courbin et al., 2019). However, waterholes remain key hunting sites for lions and some waterholes remain consistently more visited by herbivores than others (Chamaillé-Jammes et al., 2016) and may therefore be seen as sites with more predictable prey presence by lions. As demonstrated by Patin et al. (2020), predators should therefore seek to be unpredictable to optimize their hunting success, integrating some level of randomness in their movement behaviour to maximize their chances of encountering and capturing prey. Regarding this predator-prey space race, lions would therefore benefit from not visiting too regularly the same waterholes in order to be less predictable to prey. Lions in Hwange National Park are known to make most of their kills close to a waterhole and to commonly depart from a kill site to visit another waterhole (Valeix et al., 2011). Altogether, this and our findings suggest that lions may spread out their hunting grounds in an unpredictable manner. Even in the absence of kills, lions need to depart sufficiently quickly from a given hunting patch to counter-balance the anti-predator responses adopted by prey once they detect their presence there (Charnov et al. 1976; Kotler 1992). One of the anti-predator responses adopted by prey is to move away from the site of an encounter with lions (Courbin et al., 2016; Martin & Owen-Smith, 2016; Say-Sallaz et al., 2023), possibly leading to a depression in prey abundance in the vicinity of a waterhole following a lion visit. Here, unfortunately, our prey-related variables do not allow us to draw inference about these short-term variations in prey abundance as they cannot be used to predict the number of prey at waterhole at the specific time of the visit by a lion and may therefore not perfectly reflect prey abundance from a lion’s viewpoint. Indeed, although some waterholes have consistently greater herbivore abundance than others in Hwange National Park (Chamaillé-Jammes et al., 2016), lions may use other cues to infer instantaneous prey abundance. Because of a short-term depression in prey abundance, one waterhole having consistently greater herbivore abundance than others may offer – at a given time, for example following an attempted hunt – less opportunities for lions to encounter prey than an undisturbed waterhole having on average a lower number of herbivores in its vicinity. Being territorial animals (Schaller, 1972), lions have access to only a limited number of waterholes to spread out their hunting effort. Therefore, even if they seek preferentially to visit undisturbed feeding patches, where prey are most available, lions may be constrained to visit patches with low prey abundance or poor prey catchability if those patches are the ones least disturbed by their previous presence.

### A territorial and social species

Although this study focuses on the importance of prey-related variables to explain between-waterhole variations in the duration and frequency of visits to a waterhole, lions are not solely predators but also territorial and social animals (Schaller, 1972). Other variables, unrelated to prey, may therefore partly explain differences in the frequency and duration of visits to a waterhole. In particular, beyond being key hunting patches, waterholes are key marking places for lions (Dejeante, Loveridge, Macdonald, Madhlamoto, Chamaillé-Jammes, et al., 2025). For example, lions vocalize more frequentlyly near water sources (Wijers et al. 2021), and male lions more frequently encounter their rivals near waterholes (Dejeante et al. 2024). The distribution of these waterholes greatly structures the size and shape of lion territories (Valeix et al., 2012) and drives their territorial movements in the core and in the periphery of the territory (Dejeante, Loveridge, Macdonald, Madhlamoto, Chamaillé-Jammes, et al., 2025). We would therefore expect that lions’ territorial behaviours – and not solely predator-prey interactions – influence the duration and the frequency of lion visits to a waterhole. As demonstrated for another territorial species (wolves that integrate space and time to visit territorial borders; Schlägel et al. 2017), lions might, for example, be more likely to visit a waterhole not visited for a long time to renew their marking signals at this place. Interestingly, here, we found a great influence of the vegetation openness around a waterhole: male lions were more likely to frequently visit a waterhole if it was surrounded by open habitats. This result is not intuitively supportive of the prey-catchability hypothesis, but prey catchability may still matter in these large open areas, but at a finer scale than captured by our metrics. In addition, because a more open habitat is likely associated with a higher probability of encountering large herds of large herbivores in Hwange National Park, this result could be shaped by a long-term cue of high prey abundance, not captured by the data available from waterhole counts. However, other explanations are possible. In particular, a territorial-related explanation is that male lions spend more time with their pride in open areas because the visibility is greater in these habitats, allowing them to detect more easily intruders and so to less frequently patrol their territory (Dejeante et al., 2024; Funston et al., 1998). If male lions favour visiting waterhole surrounded by open habitats for territorial reasons not related to prey, these prey could, however, become reluctant to use these waterholes. This would lead to a mismatch between prey abundance and the patterns of lion visits to a waterhole. At the interspecific level, the spatio-temporal use of waterholes may also be partly driven by carnivore intraguild interactions. In the Hwange ecosystem, this is for example the case with lions often encountering spotted hyaenas around waterholes and leaving the waterhole area after such encounters (Périquet et al., 2021).

## Conclusion

Many studies investigating the role of waterholes, i.e., prey patches, in the spatial behaviour of lions implicitly assumed that all waterholes are the same (Davidson et al., 2012, 2013; Valeix et al., 2010; Valeix, Fritz, et al., 2009). Here – to our knowledge for the first time – we investigated whether this assumption was true. Clearly there is an heterogeneity in the frequency and duration of lion visits at waterholes, suggesting that not all waterholes are the same from a lion’s viewpoint. Although lions did not equally use all waterholes within their territory, we were able to provide little evidence that the variability of the frequency and duration of lion visits to a waterhole was determined by between-waterhole variations in prey abundance and our measures of its catchability. Three, non-exclusive, main points can explain this result: (1) lions and their prey are involved in a “shell-game" that leads them to adopt unpredictable movement strategies, and the proxies used in this work did not take into account these dynamics, (2) lions have access to a limited number of waterholes amongst which to distribute their hunting effort, and this leads them to visit some waterholes with lower prey abundance or catchability solely because those are the least disturbed, (3) lions do not visit waterholes solely to hunt but also to interact with conspecifics (both social mates or rivals) for example to frequently renew their territorial signals at these places, whatever the abundance and catchability of prey is there.

## Acknowledgments

We thank the Zimbabwe Parks and Wildlife Management Authority for permission to carry out this research. We are deeply grateful to Jane Hunt, Zeke Davidson, Nicholas Elliot, Brent Stapelkamp, Dan Parker, Agrippa Moyo, Lovemore Sibanda, Moreangels Mbizah, Andrea Sibanda and Liomba Mathe for their roles in the collection of lion GPS data. We are also indebted to Wildlife Environment Zimbabwe (WEZ) for their invaluable long-term dataset on waterhole use by herbivores, and we sincerely thank Yolan Richard for measuring the distances between the waterholes’ bank and the vegetation cover. Finally, we thank Kirby Mills and Andrew Kittle for their comments on an earlier version of this manuscript. Preprint version 3 of this article has been peer-reviewed and recommended by Peer Community In Ecology (https://doi.org/10.24072/pci.ecology.100802; Chaverri, 2026)

## Data Availability

Data and codes used in this study are publicly available in a figshare repository (Dejeante, Loveridge, Macdonald, Madhlamoto, Chamaillé, et al., 2025)

## Supplementary Information

**Table S1.** Results using a 2km distance threshold to define lion visits to a waterhole. Summary of the candidate models explaining the variability in the frequency of lion visits to a waterhole. The frequency of lion visits to a waterhole is calculated as the number of visits over 100 days (between early July and mid-October; i.e., during the late dry season). All models included the number of waterholes within the territory of each animal-year and a random effect with the identity of each animal-year. The most parsimonious model (ΔAICc < 2 and lowest number of parameters *K*) is shown in bold.

| Frequency of visits to a waterhole – 2km |  | Female lion |  |  | Male lion |  |  |
| --- | --- | --- | --- | --- | --- | --- | --- |
| Candidate models | K | AICc | $\Delta AICc$ | R<sup>2</sup> | AICc | $\Delta AICc$ | R<sup>2</sup> |
| *Null hypothesis* |  |  |  |  |  |  |  |
| Number of waterholes | 4 | 1939,6 | 17,7 | 0,32 | 2652,8 | 64,8 | 0,06 |
| *Prey abundance hypothesis* |  |  |  |  |  |  |  |
| Herd | 5 | 1933,9 | 12,0 | 0,35 | 2643,5 | 55,4 | 0,10 |
| Buffalo | 5 | 1937,9 | 16,0 | 0,33 | 2651,3 | 63,2 | 0,07 |
| Herd + Buffalo | 6 | 1935,0 | 13,1 | 0,35 | 2645,0 | 56,9 | 0,10 |
| *Functional response in prey-abundance hypothesis* |  |  |  |  |  |  |  |
| Herd × Absolute Herd | 7 | 1936,4 | 14,5 | 0,35 | 2644,8 | 56,7 | 0,11 |
| Buffalo × Absolute Buffalo | 7 | 1937,2 | 15,3 | 0,34 | 2653,9 | 65,8 | 0,08 |
| Herd × Absolute Herd + Buffalo × Absolute Buffalo | 10 | 1939,6 | 17,7 | 0,36 | 2649,6 | 61,6 | 0,11 |
| *Prey catchability hypothesis* |  |  |  |  |  |  |  |
| Distance to cover | 5 | 1939,1 | 17,1 | 0,32 | 2647,3 | 59,3 | 0,09 |
| Vegetation openness | 5 | 1924,5 | 2,6 | 0,35 | 2593,7 | 5,7 | 0,23 |
| Distance to cover + Vegetation openness | 6 | 1926,6 | 4,6 | 0,35 | 2594,5 | 6,4 | 0,23 |
| *Functional response in prey-catchability hypothesis* |  |  |  |  |  |  |  |
| Distance to cover × Absolute Herd | 7 | 1941,6 | 19,7 | 0,33 | 2648,9 | 60,9 | 0,10 |
| Vegetation openness × Absolute Herd | 7 | 1927,0 | 5,0 | 0,36 | 2595,0 | 6,9 | 0,23 |
| Distance to cover × Absolute Buffalo | 7 | 1942,1 | 20,2 | 0,33 | 2645,1 | 57,0 | 0,11 |
| Vegetation openness × Absolute Buffalo | 7 | 1928,0 | 6,1 | 0,35 | 2591,2 | 3,1 | 0,25 |
| Distance to cover × (Absolute Herd + Absolute Buffalo) | 9 | 1944,0 | 22,1 | 0,34 | 2645,4 | 57,3 | 0,13 |
| Vegetation openness × (Absolute Herd + Absolute Buffalo) | 9 | 1930,0 | 8,1 | 0,36 | 2591,1 | 3,0 | 0,26 |
| Vegetation openness + Distance to cover × (Absolute Herd + Absolute Buffalo) | 10 | 1930,2 | 8,3 | 0,37 | 2589,7 | 1,6 | 0,27 |
| Distance to cover + Vegetation openness × (Absolute Herd + Absolute Buffalo) | 10 | 1932,1 | 10,2 | 0,36 | 2592,4 | 4,3 | 0,26 |
| ***Prey abundance + Prey catchability hypothesis*** |  |  |  |  |  |  |  |
| Herd + Distance to cover | 6 | 1933,9 | 12,0 | 0,35 | 2640,6 | 52,5 | 0,11 |
| Herd + Vegetation openness | 6 | **1923,6** | **1,7** | **0,36** | 2593,5 | 5,4 | 0,24 |
| Herd + Distance to cover + Habitat openness | 7 | 1925,7 | 3,8 | 0,36 | 2594,3 | 6,2 | 0,24 |
| Buffalo + Distance to cover | 6 | 1938,1 | 16,1 | 0,33 | 2646,2 | 58,2 | 0,10 |
| Buffalo + Vegetation openness | 6 | **1923,6** | **1,7** | **0,36** | **2589,8** | **1,8** | **0,25** |
| Buffalo + Distance to cover + Vegetation openness | 7 | 1925,5 | 3,6 | 0,36 | 2590,2 | 2,1 | 0,25 |
| Herd + Buffalo + Distance to cover | 7 | 1935,3 | 13,4 | 0,35 | 2642,0 | 54,0 | 0,12 |
| Herd + Buffalo + Vegetation openness | 7 | 1924,3 | 2,4 | 0,36 | 2591,5 | 3,4 | 0,25 |
| Herd + Buffalo + Distance to cover + Vegetation openness | 8 | 1926,3 | 4,4 | 0,36 | 2591,9 | 3,8 | 0,25 |
| ***Prey abundance × Prey catchability hypothesis*** |  |  |  |  |  |  |  |
| Herd × Distance to cover | 7 | 1935,4 | 13,5 | 0,35 | 2637,4 | 49,3 | 0,13 |
| Herd × Habitat openness | 7 | 1921,9 | 0 | 0,37 | 2595,4 | 7,4 | 0,24 |
| Herd × (Distance to cover + Vegetation openness) | 9 | 1925,9 | 3,9 | 0,37 | 2598,1 | 10,0 | 0,24 |
| Buffalo × Distance to cover | 7 | 1939,0 | 17,1 | 0,34 | 2646,1 | 58,1 | 0,11 |
| Buffalo × Vegetation openness | 7 | 1925,5 | 3,6 | 0,35 | 2591,0 | 2,9 | 0,25 |
| Buffalo × (Distance to cover + Vegetation openness) | 9 | 1927,1 | 5,2 | 0,36 | 2590,1 | 2,0 | 0,26 |
| Herd × Distance to cover + Vegetation openness | 8 | 1927,7 | 5,7 | 0,36 | 2596,1 | 8,0 | 0,24 |
| Herd × Vegetation openness + Distance to cover | 8 | 1924,0 | 2,1 | 0,37 | 2596,1 | 8,1 | 0,24 |
| Buffalo × Distance to cover + Vegetation openness | 8 | 1925,6 | 3,6 | 0,36 | 2588,1 | 0 | 0,26 |
| Buffalo × Vegetation openness + Distance to cover | 8 | 1927,4 | 5,5 | 0,35 | 2591,4 | 3,4 | 0,25 |
| (Herd + Buffalo) × Distance to cover + Habitat openness | 10 | 1927,7 | 5,8 | 0,37 | 2590,8 | 2,7 | 0,27 |
| (Herd + Buffalo) × Vegetation openness + Distance to cover | 10 | 1926,4 | 4,5 | 0,37 | 2593,7 | 5,6 | 0,26 |
| (Herd + Buffalo) × (Vegetation openness + Distance to cover) | 12 | 1928,5 | 6,6 | 0,38 | 2594,4 | 6,3 | 0,27 |

**Table S2.** Results using a 2km distance threshold to define lion visits to a waterhole. Summary of the candidate models explaining the variability in the duration of lion visits to a waterhole. All models included the number of waterholes within the territory of each animal-year and a random effect with the identity of each animal-year. The most parsimonious model (ΔAICc < 2 and lowest number of parameters *K*) is shown in bold.

| Duration of visit to a waterhole – 2km |  | Female lion |  |  | Male lion |  |  |
| --- | --- | --- | --- | --- | --- | --- | --- |
| Candidate models | K | AICc | $\Delta\text{AICc}$ | R<sup>2</sup> | AICc | $\Delta\text{AICc}$ | R<sup>2</sup> |
| *Null hypothesis* |  |  |  |  |  |  |  |
| Number of waterholes | 4 | **4684,2** | **5,0** | **0,0** | 5742,3 | 6,5 | 0,07 |
| *Prey abundance hypothesis* |  |  |  |  |  |  |  |
| Herd | 5 | 4684,6 | 5,5 | 0,01 | 5740,2 | 4,3 | 0,08 |
| Buffalo | 5 | 4685,4 | 6,3 | 0,0 | 5744,4 | 8,5 | 0,07 |
| Herd + Buffalo | 6 | 4686,5 | 7,3 | 0,01 | 5742 | 6,1 | 0,08 |
| *Functional response in prey-abundance hypothesis* |  |  |  |  |  |  |  |
| Herd × Absolute Herd | 7 | 4684,3 | 5,2 | 0,04 | 5741,2 | 5,4 | 0,10 |
| Buffalo × Absolute Buffalo | 7 | 4687,1 | 7,9 | 0,02 | 5747,8 | 12,0 | 0,07 |
| Herd × Absolute Herd + Buffalo × Absolute Buffalo | 10 | 4689,5 | 10,3 | 0,04 | 5747,2 | 11,3 | 0,10 |
| *Prey catchability hypothesis* |  |  |  |  |  |  |  |
| Distance to cover | 5 | 4684,3 | 5,2 | 0,01 | 5743,5 | 7,6 | 0,07 |
| Vegetation openness | 5 | 4684,6 | 5,5 | 0,01 | **5736,0** | **0,1** | **0,09** |
| Distance to cover + Habitat openness | 6 | 4681,4 | 2,2 | 0,03 | 5737,7 | 1,8 | 0,09 |
| *Functional response in prey-catchability hypothesis* |  |  |  |  |  |  |  |
| Distance to cover × Absolute Herd | 7 | 4684,8 | 5,6 | 0,03 | 5744,2 | 8,3 | 0,09 |
| Vegetation openness × Absolute Herd | 7 | 4684,2 | 5,1 | 0,04 | 5737,0 | 1,1 | 0,11 |
| Distance to cover × Absolute Buffalo | 7 | 4687,0 | 7,8 | 0,01 | 5746,8 | 10,9 | 0,08 |
| Vegetation openness × Absolute Buffalo | 7 | 4683,1 | 3,9 | 0,03 | 5739,0 | 3,1 | 0,10 |
| Distance to cover × (Absolute Herd + Absolute Buffalo) | 9 | 4688,2 | 9,0 | 0,04 | 5748 | 12,1 | 0,09 |
| Vegetation openness × (Absolute Herd + Absolute Buffalo) | 9 | 4684,6 | 5,4 | 0,05 | 5740,6 | 4,7 | 0,11 |
| Vegetation openness + Distance to cover × (Absolute Herd + Absolute Buffalo) | 10 | 4684,1 | 5,0 | 0,06 | 5742,1 | 6,2 | 0,11 |
| Distance to cover + Vegetation openness × (Absolute Herd + Absolute Buffalo) | 10 | 4680,0 | 0,8 | 0,08 | 5742,4 | 6,5 | 0,11 |
| ***Prey abundance + Prey catchability hypothesis*** |  |  |  |  |  |  |  |
| Herd + Distance to cover | 6 | 4684,7 | 5,5 | 0,01 | 5741,7 | 5,8 | 0,08 |
| Herd + Vegetation openness | 6 | 4685,5 | 6,4 | 0,01 | 5735,9 | 0,0 | 0,10 |
| Herd + Distance to cover + Vegetation openness | 7 | 4682,5 | 3,4 | 0,03 | 5737,6 | 1,7 | 0,10 |
| Buffalo + Distance to cover | 6 | 4685,2 | 6,0 | 0,01 | 5745,5 | 9,6 | 0,07 |
| Buffalo + Vegetation openness | 6 | 4685,8 | 6,7 | 0,01 | 5738,0 | 2,1 | 0,09 |
| Buffalo + Distance to cover + Vegetation openness | 7 | 4681,5 | 2,3 | 0,03 | 5739,7 | 3,8 | 0,09 |
| Herd + Buffalo + Distance to cover | 7 | 4686,3 | 7,1 | 0,01 | 5743,4 | 7,5 | 0,08 |
| Herd + Buffalo + Vegetation openness | 7 | 4687,3 | 8,1 | 0,01 | 5737,9 | 2,0 | 0,10 |
| Herd + Buffalo + Distance to cover + Vegetation openness | 8 | 4683,4 | 4,3 | 0,03 | 5739,7 | 3,8 | 0,10 |
| ***Prey abundance × Prey catchability hypothesis*** |  |  |  |  |  |  |  |
| Herd × Distance to cover | 7 | 4685,9 | 6,7 | 0,02 | 5743,7 | 7,8 | 0,08 |
| Herd × Habitat openness | 7 | 4683,1 | 3,9 | 0,03 | 5737,6 | 1,7 | 0,10 |
| Herd × (Distance to cover + Vegetation openness) | 9 | 4682,7 | 3,5 | 0,04 | 5741,5 | 5,6 | 0,10 |
| Buffalo × Distance to cover | 7 | 4687,3 | 8,1 | 0,01 | 5747,6 | 11,7 | 0,07 |
| Buffalo × Vegetation openness | 7 | 4683,9 | 4,7 | 0,02 | 5739,9 | 4,0 | 0,09 |
| Buffalo × (Distance to cover + Vegetation openness) | 9 | 4681,2 | 2,0 | 0,05 | 5743,7 | 7,8 | 0,09 |
| Herd × Distance to cover + Vegetation openness | 8 | 4683,4 | 4,2 | 0,03 | 5739,6 | 3,7 | 0,10 |
| Herd × Habitat openness + Distance to cover | 8 | 4680,6 | 1,4 | 0,04 | 5739,4 | 3,5 | 0,10 |
| Buffalo × Distance to cover + Vegetation openness | 8 | 4683,6 | 4,4 | 0,03 | 5741,7 | 5,8 | 0,09 |
| Buffalo × Vegetation openness + Distance to cover | 8 | 4679,2 | 0 | 0,05 | 5741,6 | 5,7 | 0,09 |
| (Herd + Buffalo) × Distance to cover + Vegetation openness | 10 | 4686,1 | 7,0 | 0,04 | 5743,8 | 7,9 | 0,10 |
| (Herd + Buffalo) × Vegetation openness + Distance to cover | 10 | 4681,7 | 2,5 | 0,05 | 5743,5 | 7,6 | 0,10 |
| (Herd + Buffalo) × (Vegetation openness + Distance to cover) | 12 | 4685,4 | 6,2 | 0,06 | 5747,7 | 11,8 | 0,10 |

**Table S3.**
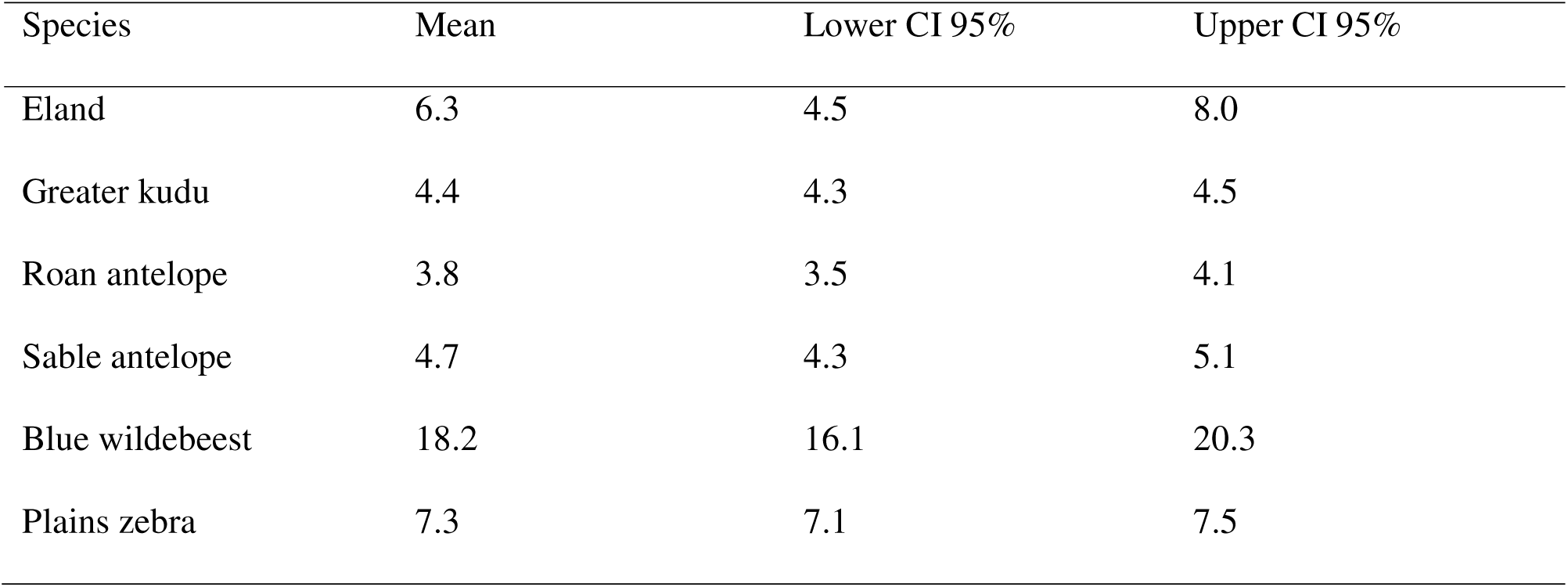
Herd size of medium-sized prey in Hwange National Park. These sizes were calculated with both adult and juvenile individuals.

| Species | Mean | Lower CI 95% | Upper CI 95% |
| --- | --- | --- | --- |
| Eland | 6.3 | 4.5 | 8.0 |
| Greater kudu | 4.4 | 4.3 | 4.5 |
| Roan antelope | 3.8 | 3.5 | 4.1 |
| Sable antelope | 4.7 | 4.3 | 5.1 |
| Blue wildebeest | 18.2 | 16.1 | 20.3 |
| Plains zebra | 7.3 | 7.1 | 7.5 |

**Table S4.** Coefficients, standard errors and p-values for the negative binomial regressions testing the influence of prey abundance and catchability on the frequency of visits to waterholes by male lions. Positive coefficients indicate for higher frequencies of visits.

| Predictor | Estimate | Std. Error | z value | p-value |
| --- | --- | --- | --- | --- |
| (Intercept) | 2.121 | 0.048 | 44.38 | < 0.001 |
| Number of waterholes | -0.263 | 0.054 | -4.86 | < 0.001 |
| Buffalo | 0.149 | 0.051 | 2.93 | < 0.05 |
| Habitat openness | 0.417 | 0.049 | 8.54 | < 0.001 |

**Table S5.** Coefficients, standard errors and p-values for the negative binomial regressions testing the influence of prey abundance and catchability on the frequency of visits to waterholes by female lions.

| Predictor | Estimate | Std. Error | z value | p-value |
| --- | --- | --- | --- | --- |
| (Intercept) | 1.773 | 0.067 | 26.27 | < 0.001 |
| Number of waterholes | -0.642 | 0.075 | -8.53 | < 0.001 |
| Herd | 0.158 | 0.053 | 2.95 | < 0.05 |
| Habitat openness | 0.114 | 0.053 | 2.17 | < 0.05 |

Positive coefficients indicate for higher frequencies of visits.

**Table S6.** Coefficients, standard errors and p-values for the negative binomial regressions testing the influence of prey abundance and catchability on the duration of visits to waterholes by female lions. Positive coefficients indicate for higher durations of visits.

| Predictor | Estimate | Std. Error | z value | p-value |
| --- | --- | --- | --- | --- |
| (Intercept) | 6.114 | 0.060 | 102.52 | < 0.001 |
| Number of waterholes | 0.155 | 0.069 | 2.25 | < 0.05 |

**Table S7.** Coefficients, standard errors and p-values for the negative binomial regressions testing the influence of prey abundance and catchability on Positive coefficients indicate for higher durations of the duration visits. of visits to waterholes by male lions.

| Predictor | Estimate | Std. Error | z value | p-value |
| --- | --- | --- | --- | --- |
| (Intercept) | 5.803 | 0.050 | 115.42 | < 0.001 |
| Number of waterholes | -0.203 | 0.058 | -3.52 | < 0.001 |
| Habitat openness | 0.110 | 0.044 | 2.51 | < 0.05 |

**Figure S1.**
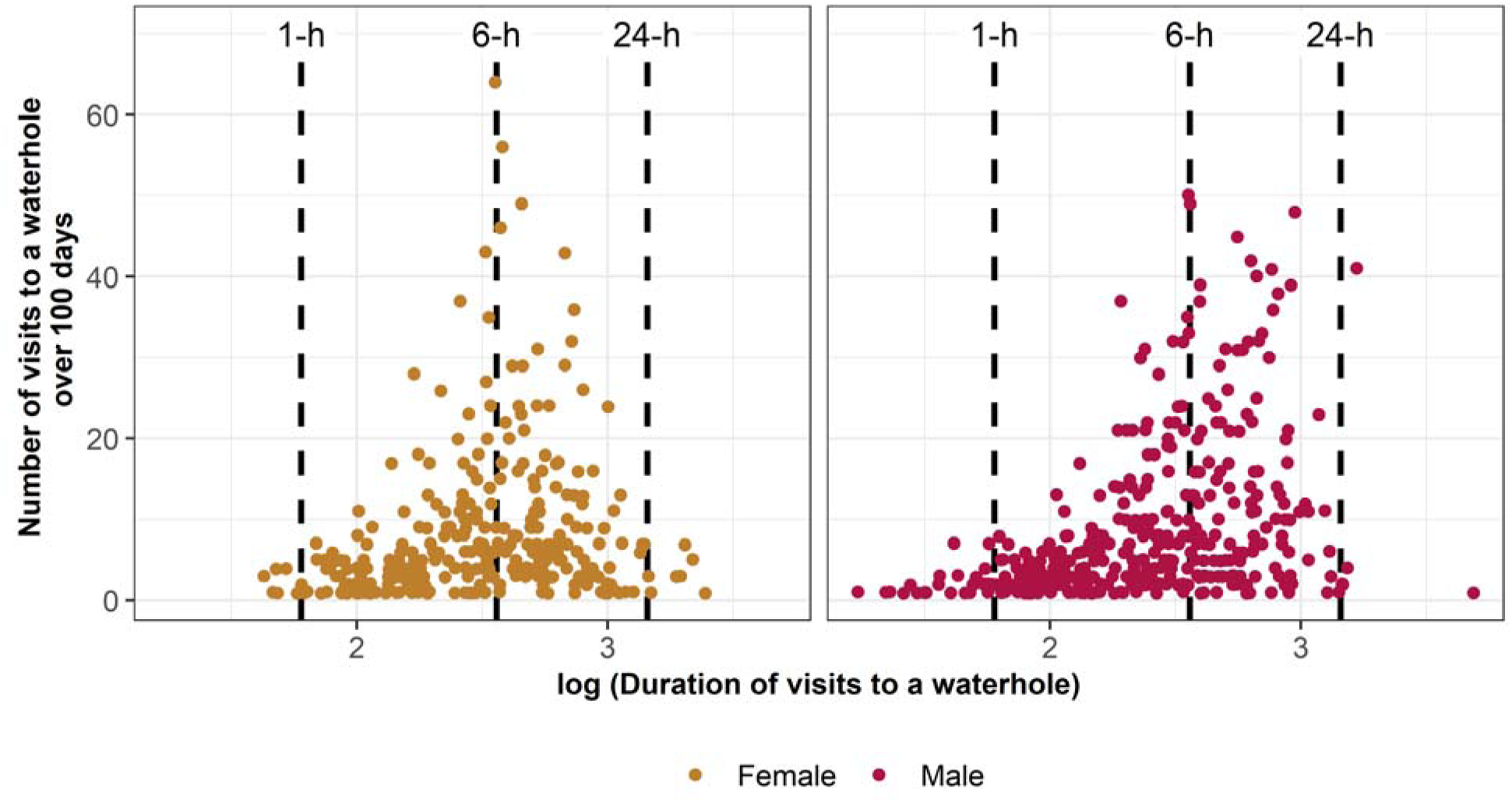
Relationship between the duration and the frequency of lion visits to a specific waterhole for females (yellow) and males (red). The duration of visits to a waterhole is in minutes but log-transform for graphical representations. Vertical dashed lines show visits to a waterhole lasting 1h, 6h and 24h.

